# A steady-state algebraic model for the time course of covalent enzyme inhibition

**DOI:** 10.1101/2020.06.10.144220

**Authors:** Petr Kuzmič

## Abstract

This report describes a double-exponential algebraic equation for the time course of irreversible enzyme inhibition following the two-step mechanism *E* + *I* ⇌ *E·I* → *EI*, under the steady-state approximation. Under the previously invoked rapid-equilibrium approximation [Kitz & Wilson (1962) *J. Biol. Chem*. **237**, 3245] it was assumed that the rate constant for the reversible dissociation of the initial noncovalent complex is very much faster than the rate constant for the irreversible inactivation step. The steady-state algebraic equation reported here removes any restrictions on the relative magnitude of microscopic rate constants. The resulting formula was used in heuristic simulations designed to test the performance of the standard rapid-equilibrium kinetic model. The results show that if the inactivation rate constant is significantly higher than the dissociation rate constant, the conventional “*k*_obs_” method is incapable of correctly distinguishing between the two-step inhibition mechanism and a simpler one-step variant, *E* + *I* → *EI*, even for inhibitors that have very high binding affinity in the reversible noncovalent step.

## 1. Introduction

The standard algebraic method of fitting irreversible inhibition data [1, Chap. 9] is based on the simplifying assumption that the reversible formation of the initial noncovalent enzyme-inhibitor complex is essentially instantaneous on the time scale of the experiment. This assumption implies that the rate constant for covalent inactivation (“*k*_inact_”) is very much smaller than the dissociation rate constant (“*k*_off_”). However, an examination of existing experimental results reveals that the typical values of rate constants for the covalent inactivation step [2, Fig. 63] are not significantly smaller than the typical dissociation rate constants of therapeutically relevant enzyme inhibitors [3]. Thus, the rapid-equilibrium approximation clearly does not hold in many experiments, in which covalent inhibitors are evaluated for potency as possible therapeutic agents. This means that the standard algebraic equations normally used to fit covalent kinetic data might not be appropriate in many cases.

One possible solution to this difficulty is to utilize mathematical models that are based on the numerical solution of systems of simultaneous first-order ordinary differential equations (ODEs). This approach avoids having to make any simplifying assumptions, either about the underlying inhibition mechanism, or about the magnitude of microscopic rate constants. For example, the software package DynaFit [4, 5], which implements a highly advanced numerical ODE solver algorithm [6], was used in the study of covalent inhibition of the EGFR kinase [7]. However, one significant disadvantage of mathematical models that rely on the numerical solution of ODE systems is that that the requisite numerical algorithms are highly complex by comparison with the closed-form algebraic equations. Most importantly, high quality ODE solving algorithms are implemented in only very few off-the-shelf software packages.

In this report we present a simple algebraic mathematical model that can be used either to simulate or to fit covalent inhibition data by using any generic software package and does not require a highly specialized ODE solving algorithm. The algebraic model presented here allows that the three microscopic rate constants for two-step covalent inhibition (“*k*_inact_”, “*k*_on_”, and “*k*_off_”) can have arbitrary values. The model is based on two simplifying assumptions. First, it is assumed that there is no inhibitor depletion, meaning that the concentration of the inhibitor is assumed to be very much higher than the concentration of the enzyme. The second simplifying assumption is that the uninhibited reaction rate is constant throughout the entire assay, meaning that the positive control progress curve can be mathematically described as a straight line.

## 2. Methods

This section describes the theoretical and mathematical methods that were used in heuristic simulation described in this report. All computations were performed by using the software package DynaFit [4, 5]. Explanation of all mathematical symbols is given in the Appendix, see *Table A.1* and *Table A.2*.

### 2.1. Kinetic mechanisms of irreversible inhibition

In this report we will consider in various contexts the kinetics mechanisms of substrate catalysis and irreversible inhibition depicted in *Figure 1*. The top reaction scheme in *Figure 1* represents the basic Michaelis-Menten reaction mechanisms for substrate catalysis [8]. In the inhibition mechanisms **A** through **C**, it is assumed that the covalent inhibitor I is kinetically competitive with the substrate S, because the inhibitor binds only to the free enzyme E and not to the Michaelis complex E·S. Kinetic mechanisms **A** and **B** both include two consecutive steps, where E·I is a reversibly formed noncovalent initial complex. However, the theoretical assumptions underlying the two kinetic models are different.

**Figure 1:**
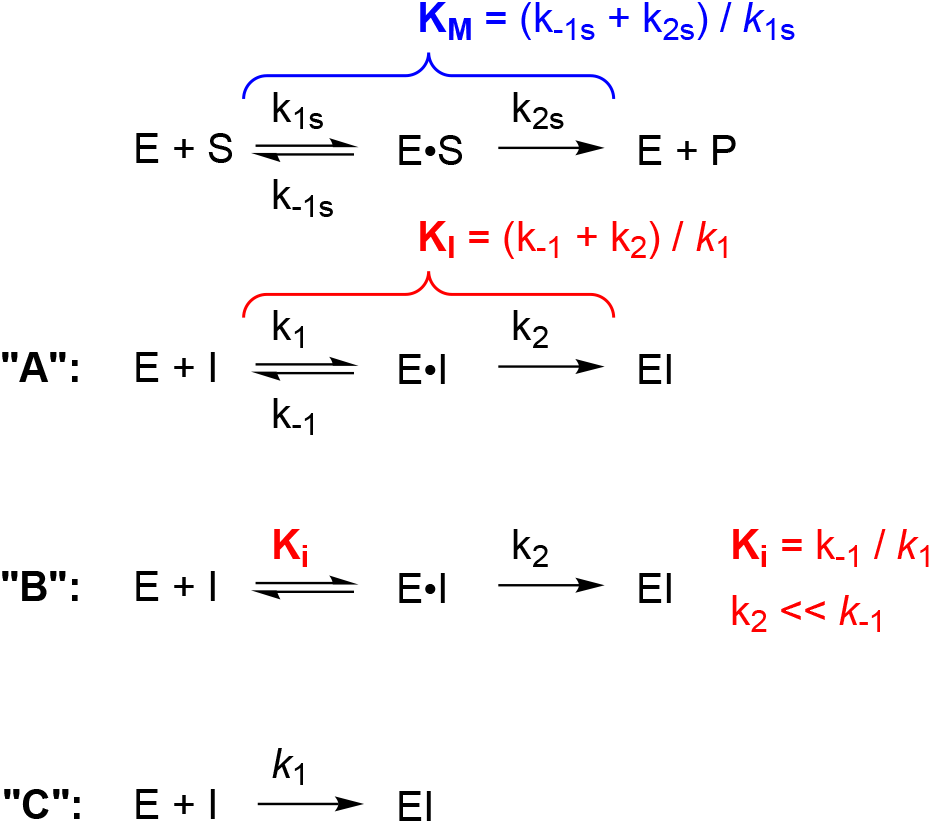
Kinetic mechanisms of substrate catalysis (top) and covalent inhibition (mechanisms **A** – **C**). For details see text.

Mechanism **A** pertains to the steady-state approximation in enzyme kinetics, where the magnitudes of the microscopic rate constants *k*_1_, *k*_−1_, and *k*_2_ can have any arbitrary values. Under those circumstances the steady-state inhibition constant is defined as *K*_I_ = (*k*_−1_ + *k*_2_)/*k*_1_ according to Malcolm & Radda [9]; the second order covalent efficiency constant also known as “*k*_inact_/*K*_i_” is defined as *k*_eff_ = *k*_1_ *k*_2_/(*k*_−1_ + *k*_2_). In contrast, mechanism **B** is invoked under the rapid equilibrium approximation [10], where it is assumed that the inactivation rate constant *k*_2_ is negligibly small compared to the dissociation rate constant *k*_−1_ and that the enzyme, inhibitor, and the noncovalent complex are always at equilibrium. If so, the steady-state inhibition constant *K*_I_ simplifies to its rapid equilibrium counterpart *K*_i_ = *k*_−1_/*k*_1_ and the second order covalent efficiency constant is defined more simply as *k*_eff_ = *k*_1_ *k*_2_/*k*_−1_.

Kinetic mechanism **C** formally describes a direct formation of the irreversibly formed covalent conjugate EI. In this case, the second-order bimolecular rate constant *k*_1_ plays the role of the covalent efficiency constant *k*_eff_. Note that under mechanism **C** there is no distinction between the steady-state and rapid-equilibrium approximations, because the noncovalent initial complex E·I is absent.

### 2.2. Mathematical models

#### 2.2.1. General mathematical model for the reaction progress

The progress of enzyme reactions is represented by Eqn (1), where *F* is the experimental signal such as fluorescence intensity; *F*_0_ is the experimental signal observed at time zero (i.e., a baseline signal as a property of the instrument); [P] is the concentration of product P at the reaction time *t* in some appropriate concentration units, such as micromoles or nanomoles per liter; and *r*_P_ is the molar response coefficient of the product under the given conditions. The molar response coefficient *r*_P_ is a proportionality constant that translates the product concentration to an experimentally observable signal, such as UV/Vis absorbance, fluorescence, or peak area, in appropriate instrument units.

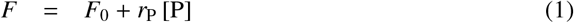

#### 2.2.2. Uninhibited substrate kinetics

In the absence of inhibitors, it is assumed that the product concentration changes over time according to the linear Eqn (4), where *v*_0_ is the uninhibited initial rate according to Eqn (2) and *t* is the reaction time. The linearity of Eqn (4) implies that the uninhibited reaction rate *v*_0_ stays effectively constant under the given experimental conditions. This in turn implies either that the initial substrate concentration is very much higher than the Michaelis constant *K*_M_ defined by Eqn (3); or that only a negligibly small fraction of the substrate S is converted to the product P at the end of the uninhibited assay; or that both of the above assumptions are satisfied.

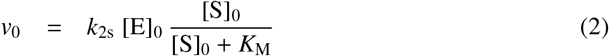

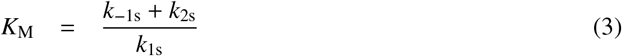

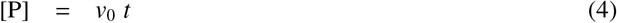

#### 2.2.3. Steady-state model for two-step covalent inhibition

In the presence of a covalent inhibitor following the steady-state mechanism **A** the concentration of product P changes over time according to the double-exponential Eqn (5). The two exponential amplitudes *a*_1_, *a*_2_ and the two first-order rate constants *r*_1_, *r*_1_ are defined by Eqns (7)–(10), respectively. The underlying assumptions are (1) zero substrate conversion, implied by the linear Eqn (4), and (2) zero inhibitor depletion, in the sense that the inhibitor concentration is very much higher than the active enzyme concentration. Exactly identical algebraic expressions for *r*_1_ and *r*_1_ were derived previously by Cornish-Bowden [11] in the context of analyzing the changes in the residual enzyme activity as opposed to the amount of reaction product.

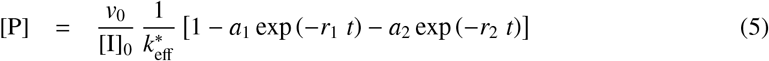

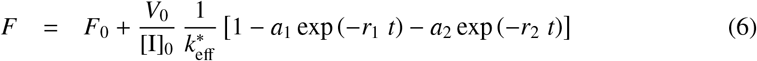

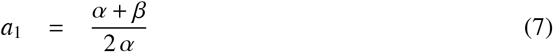

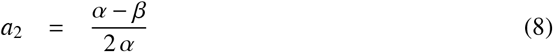

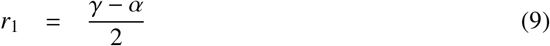

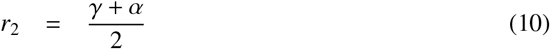

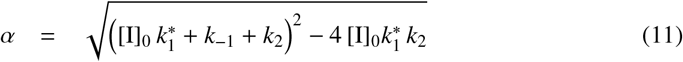

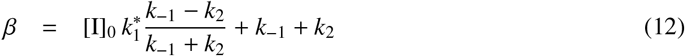

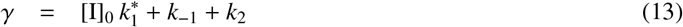

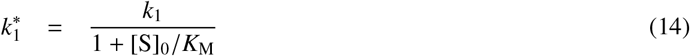

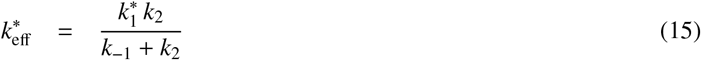

The auxiliary variables *α, β, γ*, and 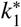 are defined by Eqns (11)–(14); *t* is the reaction time; [**I**]_0_ is the total or analytic concentration of the inhibitor, assumed to be effectively constant throughout the inhibition assay; and 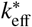 is the apparent covalent efficiency constant. Note that the sum of exponential amplitudes *a*_1_ + *a*_2_ is by definition equal to unity, because (*α* + *β*)/2*α* + (*α* − *β*)/2*α* = 1. Thus, in this sense *a*_1_ and *a*_2_ are relative amplitudes. The second exponential term, with amplitude *a*_2_, decays faster than the first term, with amplitude *a*_1_, because by definition *α* and *γ* are both positive and therefore (*α* + *γ*) > (*α* − *γ*), which implies *r*_2_ > *r*_1_ for the two first-order rate constants.

The definition of 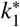 in Eqn (14) expresses the assumption that the inhibitor (I) is kinetically competitive with the substrate (S), in the sense that S and I bind to the same enzyme form, E. If the inhibitor happened to be kinetically non-competitive with the substrate, in the sense that the inhibitor would bind simultaneously and equally strongly to the free enzyme E and to the Michaelis complex E·S, the definition of 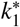 would change such that 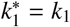. This situation could arise experimentally for example in covalent inhibition assays of protein kinases following an Ordered Bi-Bi catalytic mechanism [8], in which the inhibitor might be kinetically competitive with ATP (i.e., noncompetitive with peptide substrate) but at the same time the assay might monitor the appearance of the phosphorylated peptide, as opposed to ADP. Irreversible inhibition of bi-substrate enzymes such as protein kinases, specifically under the rapid-equilibrium approximation, is discussed in detail in ref. [12].

#### 2.2.4. Rapid-equilibrium model for two-step covalent inhibition

Under the rapid-equilibrium approximation symbolized by the kinetic mechanism **B**, the product concentration changes over time according to Eqn (16) previously derived by Tian & Tsou [13]; see their equations (A10) and (A15) in the original numbering. This algebraic model is shown here merely to highlight similarities and differences in comparison with the steady-state model represented by Eqn (5), as discussed below.

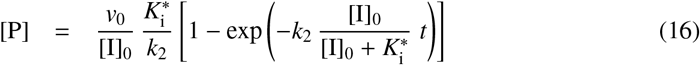

#### 2.2.5. One-step covalent inhibition

Under the simplifying assumption of (1) zero substrate conversion and (2) zero inhibitor depletion, and also assuming that the given inhibitor follows the one-step kinetic mechanism **C**, the concentration of product P changes over time according to Eqn (17). Thus, under the steadystate approximation, the two-step inhibition mechanism is described by double-exponential kinetic equation Eqn (5), whereas the one-step inhibition mechanism **C** is described by a single-exponential kinetic Eqn (17). Note that Eqn (16) was also previously derived by Tian & Tsou [13]; see their equation (A13) in the original numbering, after setting 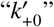 in that equation to zero, to account for purely competitive irreversible inhibition.

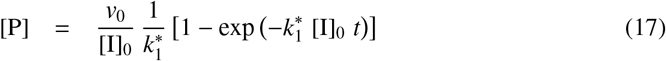

#### 2.2.6. General ODE model for two-step covalent enzyme inhibition

In the context of differential-equation modeling, the two-step inhibition mechanism **A** in *Figure 1* is mathematically represented by the ODE system defined by Eqns (18)–(24).

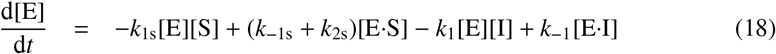

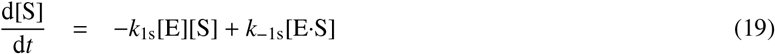

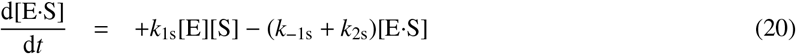

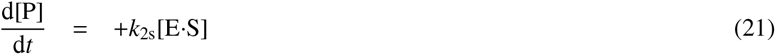

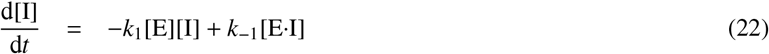

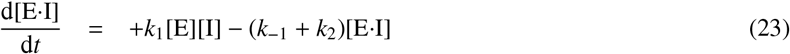

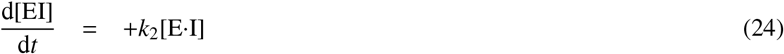

The ODE system defined by Eqns (18)–(24) was automatically generated by the software package DynaFit [5] from symbolic input. See the *Supporting Information* for details.

## 3. Results

According to Cobelli *et al*. [14], “[t]he notion of *identifiability* addresses the question of whether it is at all possible to obtain unique solutions for unknown parameters of interest in a mathematical model, from data collected in well defined stimulus-response experiments performed on a dynamic system represented by the model.” *Structural* identifiability analysis [15] is concerned with idealized data, completely free of the error, whereas *practical* identifiability analysis takes the inevitable random error into account [16]. In this section we first address both the structural and the practical identifiability of the newly derived algebraic model for covalent inhibition. We then use this model to conjure up a hypothetical irreversible inhibitor that has very strong initial binding affinity, as measured by the dissociation equilibrium constant *K*_i_ of the initial noncovalent complex, and yet apparently follows the “one-step” kinetic mechanism **C**, as if the complex **E·I** were absent.

### 3.1. Structural identifiability analysis

The results of structural identifiability analysis are illustrated in *Figure 2*. Idealized, noise-free data were simulated by using the following values of model parameters in Eqn (6): *F*_0_ = 0; *V*_0_ = 0.0005 RFU/sec;^1^ 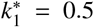 *μ*M^−1^s^−1^; *k*_−1_ = 0.001 *s*^−1^; and *k*_2_ = 0.01 s^−1^. Note that *k*_2_/*k*_−1_ = 10, meaning that that rapid-equilibrium approximation (requiring *k*_2_ ≪ *k*_−1_) does not hold. The inhibitor concentrations were 0.5, 1, 2, 4, 8, and 16 nM. The simulated time coordinates were 0, 60, 120, 180,…, 3000 seconds (50 simulated time points, stepping by one minute). The artificial data were subsequently subjected to a global fit [17] to an ODE model defined by Eqns (18)–(24). In the ODE model, the fixed parameters were [E]_0_ = 1 pM, [S]_0_ = 1 *μ*M, *k*_1s_ = 10 *μ*M^−1^s^−1^, *k*_−1s_ = 9.9 s^−1^, *k*_2s_ = 0.1 s^−1^, and *r*_P_ = 10^4^ RFU/*μ*M. Accordingly, the Michaelis constant value was *K*_M_ = (0.1 + 9.9)/10 = 1 *μ*M. The globally optimized model parameters were the molar response coefficient *r*_P_ and the rate constants *k*_1_, *k*_−1_ and *k*_2_; locally optimized^2^ model parameters were the eight baseline offsets, *F*_0_, fit separately for each progress curve. DynaFit [5] scripts that were used for the simulation and for the fitting are listed in the *Supporting Information*.

**Figure 2:**
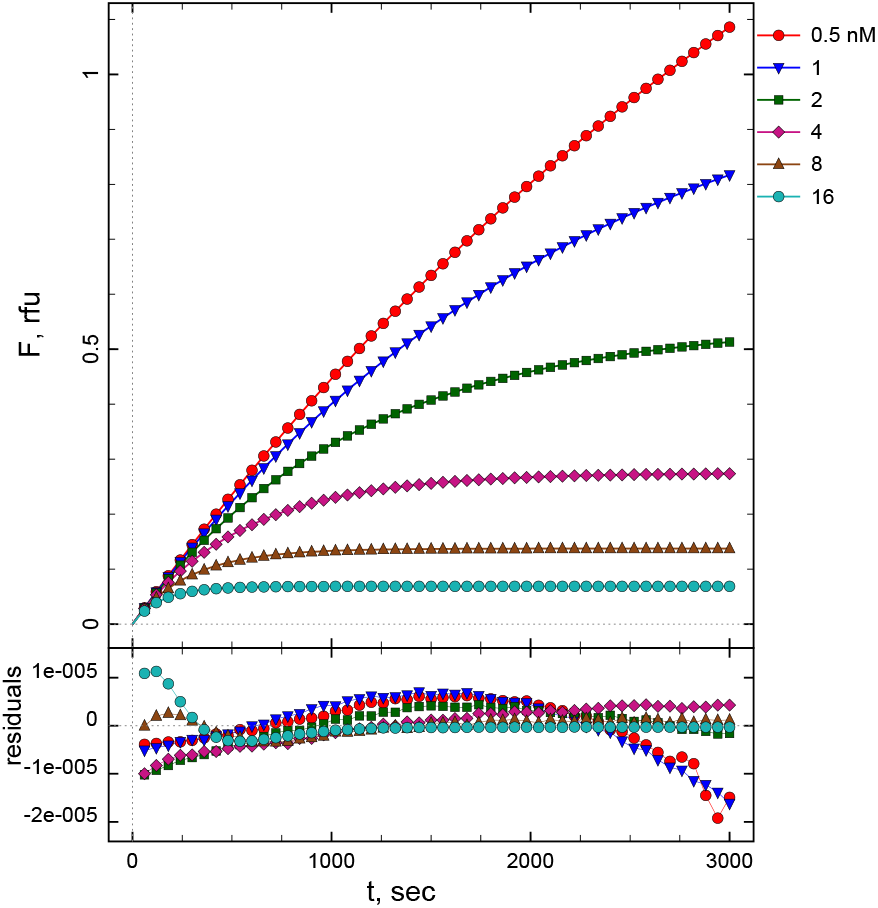
Results of structural identifiability analysis. Idealized, noise-free data were simulated by using the algebraic Eqn (5) and fit by using the ODE system Eqns (18)–(24)

The best-fit values of the globally optimized model parameters and the associated formal standard error were as follows: *k*_1_ = (1.0012 ± 0.0003) *μ*M^−1^s^−1^; *k*_−1_ = (0.00102 ± 0.00001) s^−1^; *k*_2_ = (0.01011 ±0.00003) s^−1^;and *r*_P_ = (10000.10±0.04) RFU/*μ*M. The expected best-fit value of *k*_1_ was 1 *μ*M^−1^s^−1^, because the simulated value of 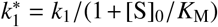 was 0.5 *μ*M^−1^s^−1^ and the adjustment factor 1 + [S]_0_/*K*_M_ in this case is equal to 1 +1/1 = 2. Thus, the fitted and theoretically expected values of all adjustable model parameters agree within five significant digits. The best-fit model curves also agree with the simulated data within five significant digits, as is illustrated in the residual plot shown as the bottom panel of *Figure 2*. The very small systematic discrepancies between the algebraic and ODE models, shown in the slightly non-random distribution of the residuals of fit, are due to the inevitable propagation of round-off and truncation errors.

The two main conclusions that can be reached from the results of this heuristic simulation study are as follows. First, the theoretical model represented by Eqn (5) is algebraically correct, because it is congruent with a fully independent mathematical representation provided by the numerical solution of an equivalent ODE system. Second, the algebraic model is structurally identifiable with respect to all three microscopic rate constants that appear in the inhibition mechanism **A**. Thus, in the purely hypothetical case of having access to entirely noise-free experimental data, it would always be possible to determine all three rate constants *k*_1_, *k*_−1_ and *k*_2_ by performing a global least-squares fit of combined reaction progress curves similar to those shown in *Figure 2*.

### 3.2. Practical identifiability analysis

In the practical identifiability study the roles of the algebraic vs. numerical models were reversed. Artificial data were simulated according to the differential-equation system Eqns (18)–(24), this time with a finite level of experimental noise added to the results. The pseudo-experimental data were subsequently fit by using the algebraic model represented by Eqn (6). The requisite DynaFit [5] input scripts are listed in the *Supporting Information*.

The simulated concentration plot in the bottom panel of *Figure 3* shows that at [**I**]_0_ = 80 nM the noncovalent complex E·I has a significant presence in the evolving reaction mixture and that this noncovalent complex is formed gradually, as opposed to instantaneously, on the time scale of the simulated experiment. These observations suggest that there is a good chance of successfully determining all three rate constants that appear in the two-step inhibition mechanism **A**.

**Figure 3:**
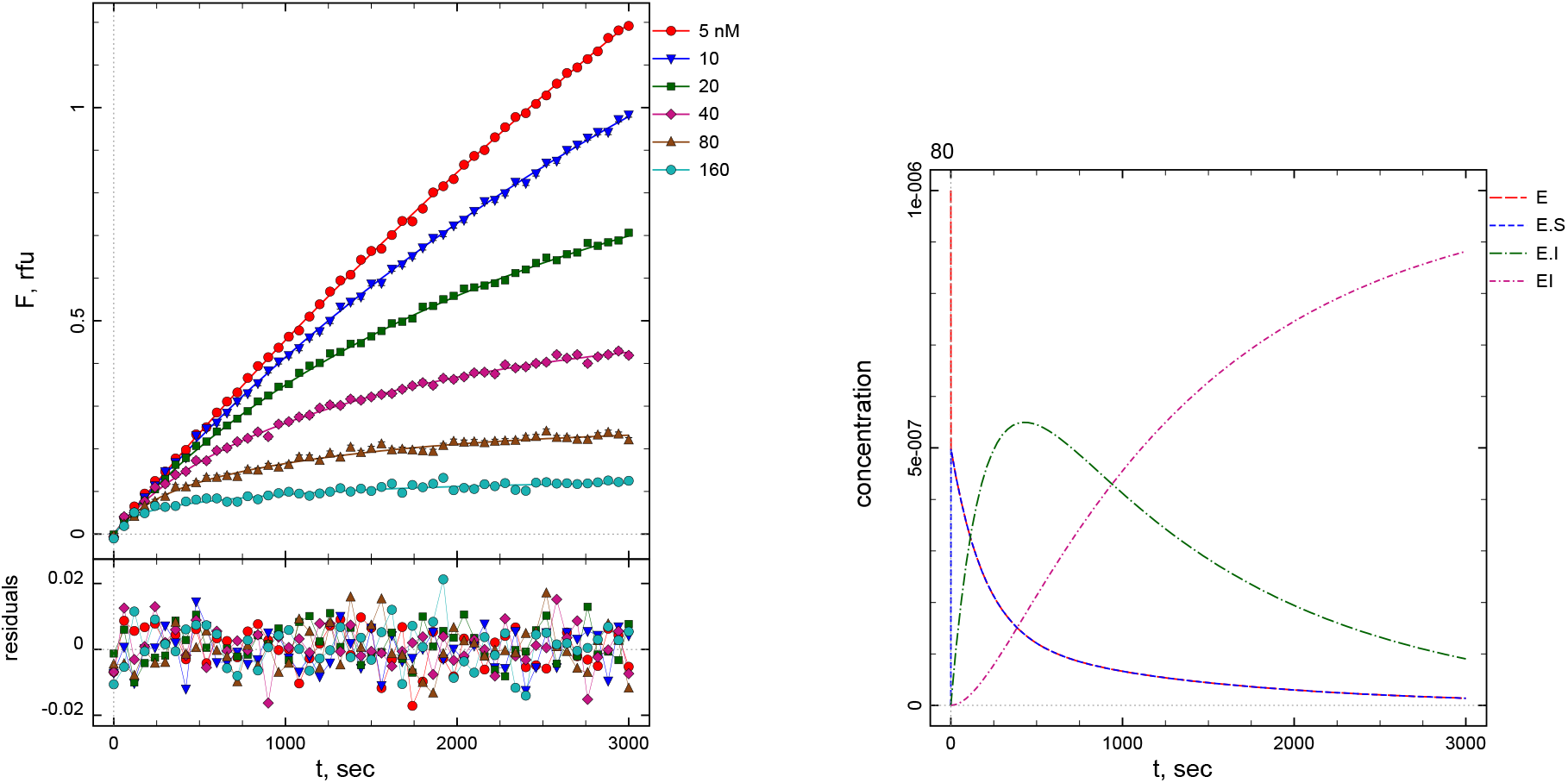
Artificial data simulated by using the ODE system Eqns (18)–(24). **Left:** Pseudo-experimental data (symbols) and the corresponding idealized model curves. **Right:** Enzyme species concentrations evolving over time at [**I**]_0_ = 80 nM. The “concentration” axis is in *μ*M units.

Simulated data shown in *Figure 3* were globally fit to the algebraic model represented by Eqn (6). The overlay of the best-fit model curves on the simulated data points was visually indistinguishable from the display of the simulated shown in *Figure 3*. The residual plots were distributed randomly, similar to the residual plot shown in *Figure 3*, upper panel. The best-fit values of globally adjustable model parameters are listed in *Table 1*. The columns labeled “low” and “high” are lower and upper limits, respectively, of non-symmetrical confidence intervals obtained by the profile-*t* method [18, 19] at 5% ΔSSQ level according to the empirical cut-off criterion advocated by Johnson [20, 21].

**Table 1:** Results of practical identifiability analysis. For details see text.

| parameter | best-fit ± std. err. | low | high |
| --- | --- | --- | --- |
| $V_0$, RFU/s | 0.0005011 ± 0.000002 | | |
| $k_1^*$, $\mu\text{M}^{-1}\text{s}^{-1}$ | 0.046 ± 0.002 | 0.040 | 0.056 |
| $k_{-1}$, $\text{s}^{-1}$ | 0.0007 ± 0.0001 | 0.0004 | 0.0014 |
| $k_2$, $\text{s}^{-1}$ | 0.0008 ± 0.0001 | 0.0003 | 0.0012 |

The results displayed in *Table 1* show that all three microscopic rate constants appearing in the steady-state two-step covalent inhibition mechanism **A** could be reliably determined from the simulated data. The non-symmetrical confidence interval for the apparent association rate constant 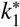 spanned from 0.046 to 0.056 *μ*M, while the theoretically expected value of 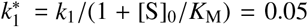 *μ*M. Similarly, the confidence intervals for *k*_−1_ and *k*_2_ were relatively narrow (high/low ratios were approximately equal to 4) and encompassed the simulated values (0.001 s^−1^ in both cases). The best-fit values of *k*_−1_ and *k*_2_ were only 20% to 30% lower than the simulated values. The main conclusion is that, for at least some combinations of microscopic rate constants appearing in the two-step mechanism **A**, all three rate constants (*k*_1_, *k*_−1_, and *k*_2_) can be determined in ordinary kinetic measurements, such as those that are typical for plate-reader assays usually performed in preclinical inhibitor screening.

### 3.3. “One-step” kinetics of a high-affinity inhibitor

Pseudo-experimental data were simulated by using the following values of model parameters in Eqn (6): *F*_0_ = 0, *V*_0_ = 0.0005 RFU/sec; 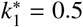 *μ*M^−1^s^−1^; *k*_−1_ = 0.001 s^−1^; and *k*_2_ = 0.01 s^−1^. Note that those are the same parameters that were used in the structural identifiability analysis, as described in section 3.1. However, in this case the simulated signal was perturbed by adding a Gaussian-distributed random noise with the standard deviation equal to 0.5% of the maximum simulated value. Each of the simulated progress curves were fit individually and separately to the standard algebraic model [1, sect. 9.1] for the time course of covalent inhibition, represented by the exponential Eqn (25), where *V*_i_ is the initial reaction rate in instrument units and *k*_obs_ is the apparent first-order rate constant corresponding to each inhibitor concentration.

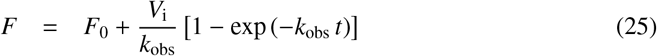

The results of fit are shown graphically in *Figure 4*. Note that the residuals of fit in the bottom panel are distributed randomly, which means that the *single-exponential* Eqn (25) is an adequate fitting model for this data, even though the data were simulated on the basis of a *double-exponential* Eqn (5). The best-fit values of the apparent first-order rate constant *k*_obs_ obtained at each inhibitor concentration are collected in *Table 2*.

**Figure 4:**
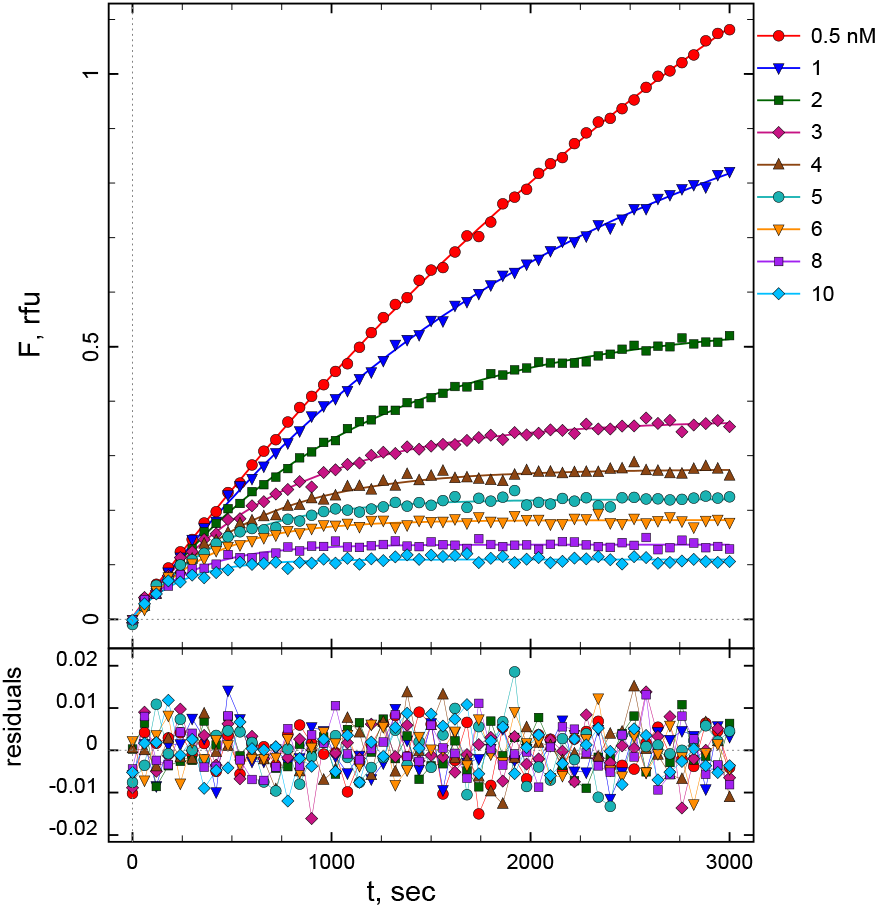
Results of fit of each individual progress curve simulated by using Eqn (5), with parameter values listed in the text, to Eqn (25), in order to determine the *k*_obs_ values associated with each inhibitor concentration listed in the right margin.

**Table 2:**
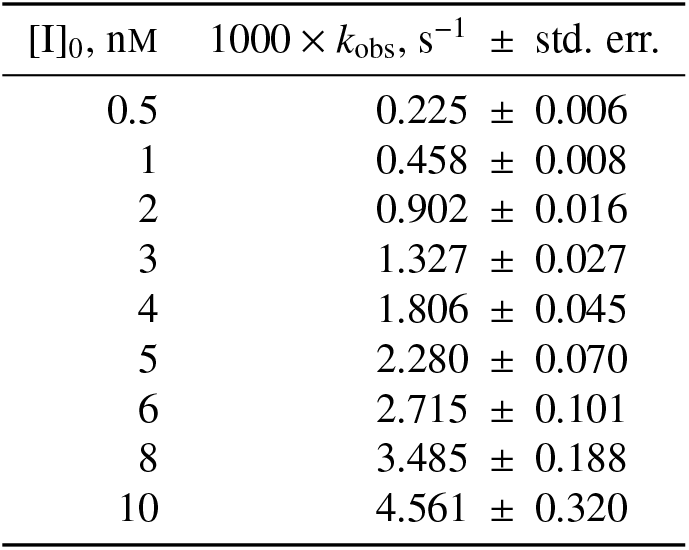
Best-fit values of the apparent first-order rate constant *k*_obs_ obtained by fitting the progress curves shown in *Figure 4* to the exponential Eqn (25).

The *k*_obs_ results collected in *Table 2* were subjected to a model discrimination analysis according to the procedure described in ref. [22], considering Eqn (26) and Eqn (27) as the two candidate fitting models, according to the standard treatment described in ref. [1, sect. 9.1] and elsewhere. The hyperbolic Eqn (26), corresponding to the two-step mechanism **B**, could be reliably excluded from consideration, using four independent statistical model selection criteria [22]. The preferred model was the linear Eqn (27), corresponding to the one-step mechanism **C**. The fit to the linear Eqn (27) produced a very well defined value of 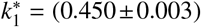 *μ*M^−1^ s^−1^, with the 95% confidence interval computed by the profile-*t* method [18,19] spanning from 0.443 to 0.457 *μ*M^−1^s^−1^. The results of fit to the linear model regression model are shown in *Figure 5*.

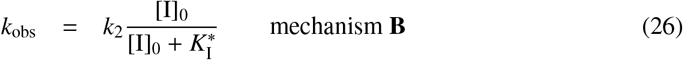

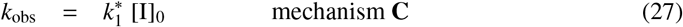

**Figure 5:**
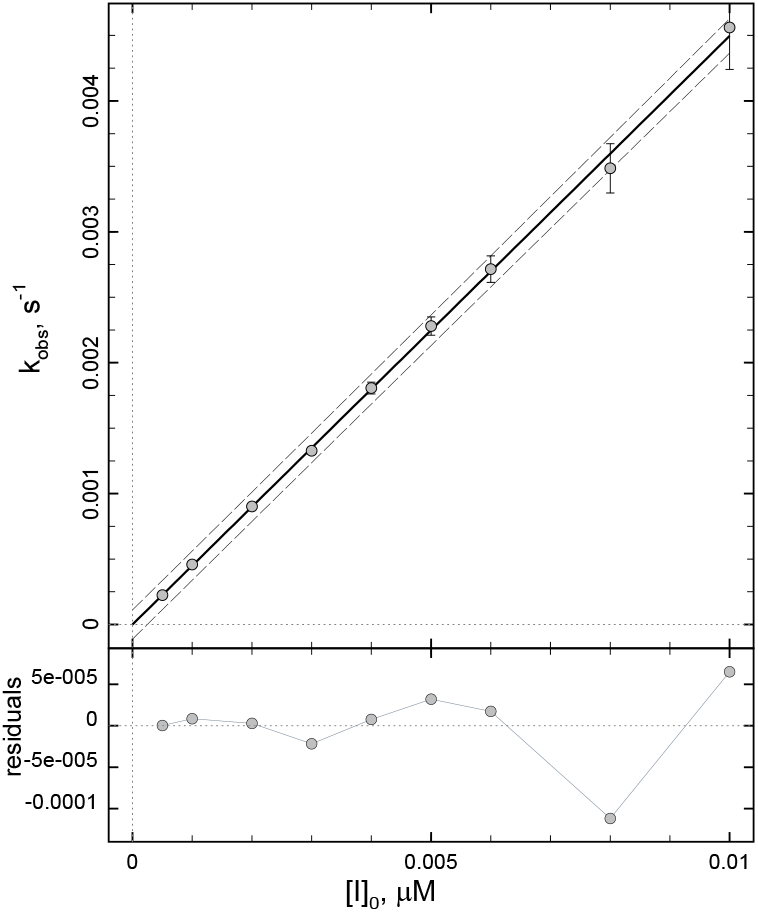
Results of fit of the *k*_obs_ values listed in *Table 2* to Eqn (27), corresponding to the one-step covalent inhibition mechanism **C**. For further details see text.

The values of microscopic rate constants used in simulating the artificial data shown in *Figure 4* were 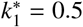 *μ*M^−1^s^−1^, *k*_−1_ = 0.001 s^−1^ and *k*_2_ = 0.01 s^−1^. The apparent covalent efficiency constant is defined as 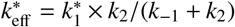. Accordingly, the “true” value of the covalent efficiency constant was 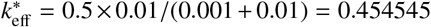 *μ*M^−1^s^−1^. Thus the best-fit value 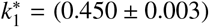 *μ*M^−1^s^−1^ and the “true” value *k*_eff_ = 0.454 *μ*M^−1^s^−1^ are in good agreement. Importantly, this agreement between the “true” and best-fit values of 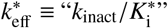 holds even though the individual values of *k*_inact_ and 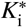 could not be determined from the simulated data.

## 4. Discussion

### Challenges in evaluating covalent inhibitors as potential drugs

Reliable evaluation of irreversible enzyme inhibitors as potential therapeutic agents is exceptionally challenging for the biochemical data analyst engaged in drug discovery. The reason is that the overall potency of covalent inhibitors consists of two separate and yet intertwined contributions. First, the inhibitor’s binding affinity is measured by the inhibition constant, *K*_i_. Second, the inhibitor’s chemical reactivity is measured by the inactivation rate constant *k*_inact_. However, we have previously documented that at least some inhibitors currently being prescribed as experimental anti-cancer drugs effectively follow the one-step kinetic mechanism **C** [22]. In those cases, the only available measure of potency is the efficiency constant *k*_eff_, also known as “*k*_inact_/*K*_i_”, which blends together both affinity and reactivity such that those two distinct molecular properties can no longer be evaluated separately.

Irreversible inhibitors express their potency in a dynamic fashion, in the sense that the residual enzyme activity decreases over time, along with the gradual evolution of the permanent covalent bond between the enzyme and the inhibitor. Thus, in order to evaluate the potency of covalent inhibitors, we need mathematical models that describe the gradual formation of the final reaction product while appropriately taking into account that the rate of product formation (i.e., enzyme activity) inevitably decreases over time. As a result, all mathematical models for the progress of covalent inhibition assays are by definition nonlinear. This complexity presents an additional challenge when compared with measuring the potency of noncovalent inhibitors, where the assay in many cases can be arranged such that the reaction progress is nearly linear.

### Existing models for the progress of covalent inhibition

Currently existing nonlinear regression models for the progress of covalent inhibition assays can be divided into two categories, according to the mathematical formalism involved. In the first category are highly advanced *differential* equation models, which eliminate any simplifying assumptions about the relative magnitude of microscopic rate constants, such as the rapid-equilibrium approximation [8]; about the relative concentration of reactants, such as requiring a very large excess of inhibitor over enzyme; or about the reaction conditions, such as assuming strict linearity of the positive control progress curve. One important disadvantage of these ODE models is that they require highly specialized software algorithms for the numerical (i.e., iterative) solution of ODE systems.

In the second category of mathematical models are *algebraic* equations, such as Eqn (25) originally derived by Tian & Tsou [13], where the definition of *k*_obs_ is given by Eqns (26)–(27). Eqn (25) applies only to covalent inhibition assays where the uninhibited positive control reaction proceeds at a strictly constant rate, meaning that the plot of product concentration over time is linear. We have previously described a closely related algebraic model that allows for the control assay to be exponential, as opposed to linear [23]. On the one hand, these algebraic models have the major advantage that they can be implemented in any software system that allows the investigator so specify an arbitrary algebraic equation. On the other hand, both of these algebraic equations are based on the *rapid-equilibrium* approximation in enzyme kinetics [8]. Accordingly, it is assumed the chemical inactivation step (rate constant *k*_2_ in *Figure 1*) is very much slower than the dissociation of the noncovalent complex (rate constant *k*_−1_).

In the specific case of the single-exponential model represented by Eqn (25), Cornish-Bowden [24, sec. 7.2.2] pointed out that “if *k*_2_ is not small enough to allow formation of E·I to be treated as an equilibrium […] the loss of activity does not follow simple first-order kinetics: there is no exact analytical solution, but the kinetics may still be analyzed by numerical methods.” However, Cornish-Bowden’s statement that there is no “analytical solution” happens to be incorrect, because the desired exact analytic (i.e., algebraic) solution does exist and is represented by the newly derived double-exponential Eqn (5).

### A need for the newly derived steady-state algebraic model

Why is it important to have at our disposal a closed-form algebraic model for the timecourse of covalent inhibition assays under the steady-state approximation? There are at least two important reasons, which are now addressed in their turn.

The first and most important need for the algebraic model newly presented in this report is that the overly simple rapid-equilibrium approximation (*k*_2_ ≪ *k*_−1_) is almost certainly violated in many practically relevant cases. This is especially true for the most promising drug candidates. The *Kinetics for Drug Discovery* (K4DD) project [3] revealed that the dissociation rate constants for therapeutically relevant enzyme inhibitors are frequently in the range corresponding to hourlong drug–target residence times, which implies *k*_−1_ < 0.0001 s^−1^ or even *k*_−1_ < 0.00001 s^−1^ in many cases. On the other hand, Abdeldayem *et al*. [2] reported that the large majority of covalent inhibitor drugs and drug candidates are associated with inactivation rate constants in the range from approximately *k*_2_ = 0.0001 s^−1^ to *k*_2_ = 0.01 s^−1^. Therefore, assuming that the initial (noncovalent, reversible) binding affinity of covalent inhibitors is reasonably similar to the binding affinity of their noncovalent structural analogs, we can conclude that the typical values of dissociation rate constants are *not* very much larger than the covalent inactivation rate constants, as is required by the rapid-equilibrium approximation. Thus, the single-exponential equation Eqn (25) for the reaction progress is very likely to be invalid, especially in the case of highly potent covalent drugs.

The second reason to have available an algebraic equation as the theoretical model for the progress of steady-state covalent inhibition assays is convenience and portability. The newly derived algebraic Eqn (5) can be implemented even in general-purpose software systems such as in Microsoft Excel, as opposed to requiring highly specialized ODE solving algorithms that are only available in very few software packages. In fact, a relevant Microsoft Excel template file is attached as one of the *Supporting Information* documents.

### “One-step” kinetics of high-affinity covalent inhibitors

One of the principal motivations for addressing the fundamental difference between the steady-state approximation and the rapid-equilibrium approximation in the analysis of covalent inhibition data has been the puzzling observation that many highly potent covalent inhibitors apparently follow the one-step kinetic mechanism **C**. The inhibition of certain protein kinases by ibrutinib represents a typical example [22, 25]. The occurrence of the one-step mechanism has been described as “nonspecific affinity labeling” in textbook literature. Small-molecule inhibitors similar to iodoacetate and N-ethyl maleimide are assumed to simultaneously modify many side-chains on the target protein molecule and have negligibly low initial binding affinity, which presumably explains their one-step kinetic behavior. In contrast, highly specific inhibitors that precisely target the enzyme’s active site are assumed to always follow the two-step kinetic mechanism **A** or **B** [24, sec. 7.2.1] [1, sec 9.1]. In this sense, the fact that certainly highly specific and high-affinity inhibitors also follow the one-step mechanism **C** might appear as a paradox.

In order to better understand the unexpected “one-step” kinetics of certain high-affinity inhibitors, in section 3.3 of this report we have conjured up an inhibitor with molecular properties that were perfectly known (i.e., simulated) in advance. The objective was to simulate a compound that might approximate the kinetic properties of ibrutinib and similar “one-step” inhibitors of protein kinases. This hypothetical inhibitor was characterized by high initial binding affinity, with inhibition constant equal to *K*_i_ = 1 nM. We have simulated assays of this inhibitor at concentrations as high as 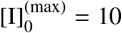 nM, which is ten times higher than the inhibition constant. Under the rapid-equilibrium approximation, it should be easily possible to determine the inhibition constant from the simulated data, because at a ten-fold excess of the inhibitor concentration over the inhibition constant the plot of *k*_obs_ vs. [I]_0_ is expected to be highly hyperbolic. The maximum observed *k*_obs_ value should be closely approaching the asymptotically saturating value, which is by definition equal to *k*_inact_. However, the actually observed *k*_obs_ vs. [I]_0_ plot was essentially linear, showing no sign of hyperbolic saturation. This means that only the covalent efficiency constant *k*_eff_ = “*k*_inact_/*K*_i_” could be determined from the simulated data, but not the values of *k*_inact_ and *K*_i_ considered separately.

The results of this simulation study confirm what has been observed experimentally for example for ibrutinib inhibition of the TEC and BTK kinases [22, 25]. A highly specific, precisely targeted, and high-affinity irreversible inhibitor characterized by an extremely low *equilibrium* dissociation constant of the reversibly formed noncovalent complex, can indeed behave *kinetically* though the reaction were proceeding via the simple one-step mechanism **C**. There are at least two possible non-mathematical explanations of this potentially puzzling behavior.

The first intuitively accessible explanation of “one-step” kinetics has to do with the familiar idea that those reversible (non-covalent) inhibitors that are characterized by extremely low dissociation rate constant *k*_−1_ can behave as effectively irreversible on the time scale of the given kinetic experiment. In fact, the distinction between truly irreversible (covalent) inhibition and effectively irreversible (non-covalent) inhibition can be so blurred that in many cases it can only be established by specialized experiments [1, sec. 5.2]. See also our previous results on extremely potent, non-covalent but nearly irreversible inhibitors of 5-α-ketosteroid reductase [26]. Importantly, if a given covalent inhibitor happens to be effectively or nearly irreversible already in the first noncovalent binding step, the overall two-step covalent inhibition mechanism will kinetically manifest as a one-step process. This is because the decrease in enzymatic activity over time is exactly identical for the one-step irreversible mechanism, *E* + *I* → *EI*, and for the two-step inhibition mechanism where both steps are irreversible, *E* + *I* → *E*·*I* → *EI*. See Appendix C for mathematical details.

Another intuitively understandable reason for the puzzling “one-step” kinetic behavior of certainly highly specific and precisely targeted irreversible inhibitors has to do with the distinction between the inhibition constant *K*_I_ and the equilibrium dissociation constant *K*_i_ of the noncovalent enzyme–inhibitor complex. As an illustrative example, consider the hypothetical case of a fully reversible inhibitor characterized by *k*_1_ = 10^6^ M^−1^s^−1^ and *k*_−1_ = 0.0001 s^−1^, corresponding to drug-target residence time of almost 3 hours. The corresponding equilibrium dissociation constant is *K*_i_ = *k*_−1_/*k*_1_ = 0.0001/1.0 = 0.0001 *μ*M = 0.1 nM. Now let us assume for the sake of discussion that the initial noncovalent binding affinity of this molecule does not deteriorate by appending to it a chemically reactive covalent warhead such that the inactivation rate constant is *k*_2_ = 0.1 s^−1^, similar to a number known cases [2]. If so, the experimentally observable inhibition constant now increases thousand-fold, because *K*_I_ = (*k*_−1_ + *k*_2_)/*k*_1_ = (0.0001 + 0.1)/1 = 0.1001 *μ*M = 100.1 nM. In a way, it is as though simply by attaching a reactive moiety to the molecule, we had somehow turned a strong binder (*K*_i_ = 0.1 nM) to a moderate-to-weak binder (*K*_I_ = 100.1 nM) without actually changing the intrinsic non-covalent binding affinity. In practical terms, it means that it is essentially impossible to sufficiently saturate the enzyme with a highly reactive covalent inhibitor under experimental conditions most commonly utilized for the evaluation of inhibitory potency. Indeed, it could easily happen that all practically useful inhibitor concentrations (i.e., those that are sufficiently low such that complete covalent inhibition will not be essentially instantaneous) might be very much lower than *K*_I_ – even as they might be very much higher than *K*_i_. If so, the compound in question will appear as a “one-step” inhibitor according to the rule-of thumb (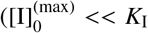 leads to apparently “one-step” kinetics) first formulated by Kitz & Wilson [10].

In conclusion, the steady-state algebraic model for two-step irreversible inhibition newly derived in this report should serve as a convenient tool to help increase our understanding of complex data-analytic issues arising in the evaluation of covalent enzyme inhibitors as potential therapeutic agents.

## Supporting information

Excel Simulation Worksheet

## Supporting information

The following supporting files accompany this document:

1. File BioKinPub-2020-03-SI.pdf: Listing of all DynaFit input scripts that were used to generate this report; instructions for using the Microsoft Excel simulation file BioKin-TN-2020-02-SI2.xls.
2. File BioKinPub-2020-03-SI2.xls: A Microsoft Excel template file that can be used to simulate the reaction progress curves according to the steady-state kinetic mechanism **A** in *Figure 1*.

## Appendix

## A. Explanation of symbols

**Table A.1:** Explanation of symbols: Microscopic rate constants and derived kinetic constants.

| Symbol | Unit | Explanation |
| --- | --- | --- |
| $k_{1s}$ | $M^{-1}s^{-1}$ | association rate constant for $E+S \rightarrow E \cdot S$ |
| $k_{-1s}$ | $s^{-1}$ | dissociation rate constant for $E \cdot S \rightarrow E + S$ |
| $k_{2s}$ | $s^{-1}$ | turnover number; $k_{2s} \equiv k_{cat}$ |
| $K_M$ | $M$ | Michaelis constant; $K_M = (k_{-1s} + k_{2s})/k_1$ |
| $k_S$ | $M^{-1}s^{-1}$ | catalytic efficiency constant; specificity constant; $k_S \equiv k_{cat}/K_M$ |
| $k_1$ | $M^{-1}s^{-1}$ | association rate constant for $E+I \rightarrow \dots$ |
| $k_{-1}$ | $s^{-1}$ | dissociation rate constant for $E \cdot I \rightarrow E + I$ |
| $k_2$ | $s^{-1}$ | inactivation rate constant; $k_2 \equiv k_{inact}$ |
| $k_1^*$ | $M^{-1}s^{-1}$ | apparent association rate constant:<br>competitive: $k_1^* = k_1/(1 + [S]_0/K_M)$<br>uncompetitive: $k_1^* = k_1(1 + [S]_0/K_M)$<br>noncompetitive $k_1^* = k_1$ |
| $k_{eff}$ | $M^{-1}s^{-1}$ | second-order inhibition efficiency constant; $k_{eff} \equiv "k_{inact}/K_i"$:<br>steady-state two-step mechanism **A**: $k_{eff} = k_1 k_2/(k_{-1} + k_2)$<br>rapid-equilibrium two-step mechanism **B**: $k_{eff} = k_1 k_2/k_{-1}$<br>one-step mechanism **C**: $k_{eff} = k_1$ |
| $k_{eff}^*$ | $M^{-1}s^{-1}$ | apparent inhibition efficiency constant:<br>competitive: $k_{eff}^* = k_{eff}/(1 + [S]_0/K_M)$<br>uncompetitive: $k_{eff}^* = k_{eff}(1 + [S]_0/K_M)$<br>noncompetitive $k_{eff}^* = k_{eff}$ |
| $K_i$ | $M$ | equilibrium dissociation constant of the $E \cdot I$ complex; $K_i = k_{-1}/k_1$ |
| $K_i^*$ | $M$ | apparent equilibrium dissociation constant:<br>competitive: $K_i^* = K_i(1 + [S]_0/K_M)$<br>uncompetitive: $K_i^* = K_i/(1 + [S]_0/K_M)$<br>noncompetitive $K_i^* = K_i$ |
| $K_I$ | $M$ | inhibition constant; $K_I = (k_{-1} + k_2)/k_1$ |
| $K_I^*$ | $M$ | apparent inhibition constant:<br>competitive: $K_I^* = K_I(1 + [S]_0/K_M)$<br>uncompetitive: $K_I^* = K_I/(1 + [S]_0/K_M)$<br>noncompetitive $K_I^* = K_I$ |
| $k_{obs}$ | $s^{-1}$ | apparent first-order rate constant for enzyme inactivation |

## B. Derivation of Eqn (5)

The derivation of the steady-state algebraic model for covalent inhibition kinetics follows the general principles utilized in ref. [27]. Accordingly, the reaction scheme displayed as mechanism **A** in the main manuscript leads to the system of Eqns (B.1)–(B.7),

**Table A.2:**
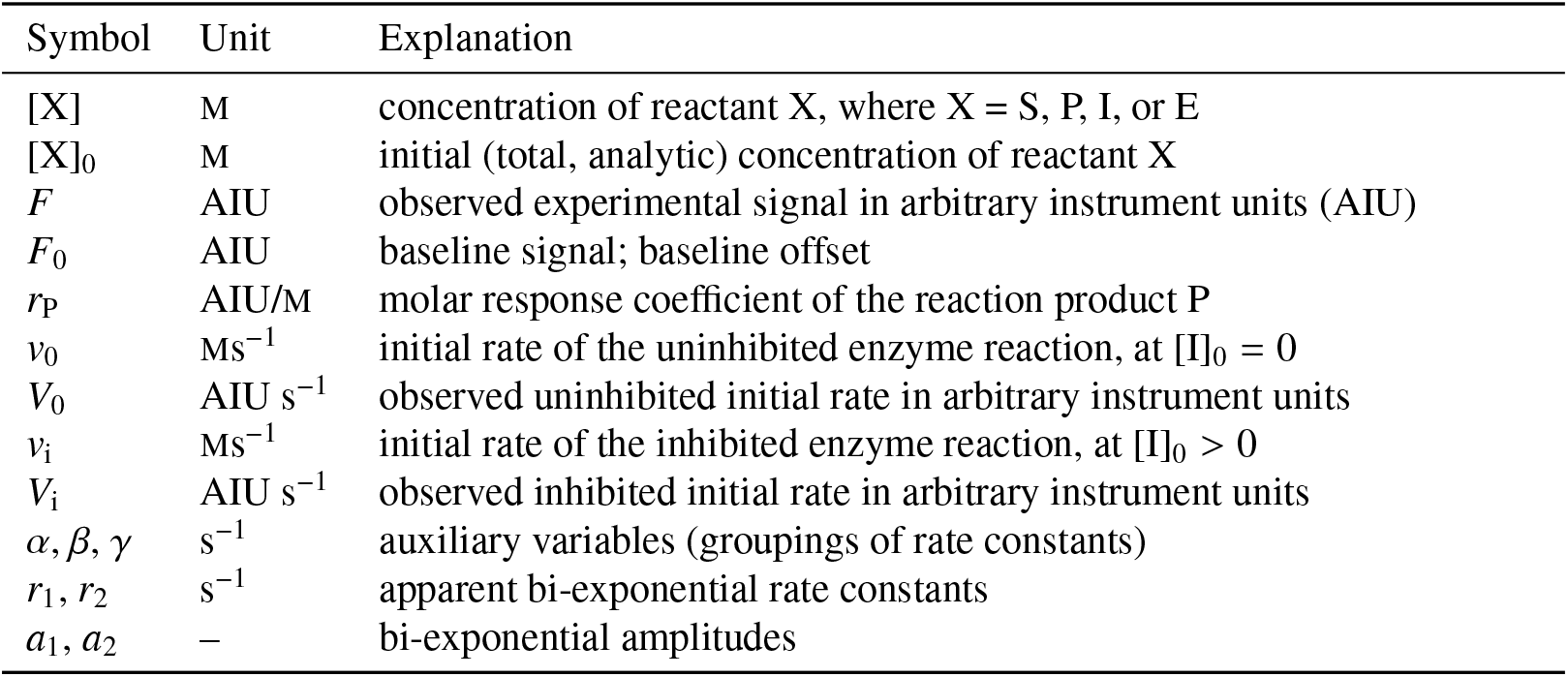
Explanation of symbols: Concentrations, reaction rates, and auxiliary symbols.

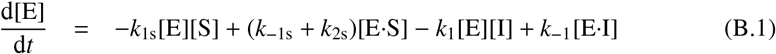

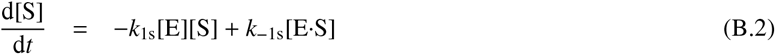

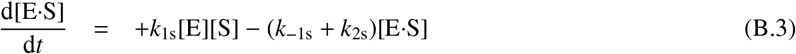

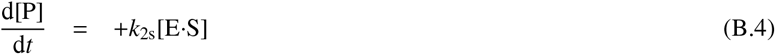

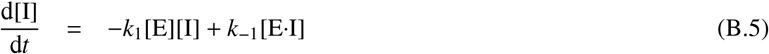

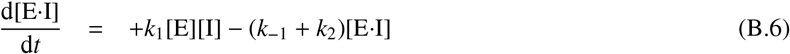

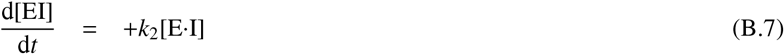

Assuming no substrate or inhibitor depletion ([I] = [I]_0_ and [S] = [S]_0_, where lower index zero represents the total or analytic concentration), we can eliminate differential equations for [I] and [S]:

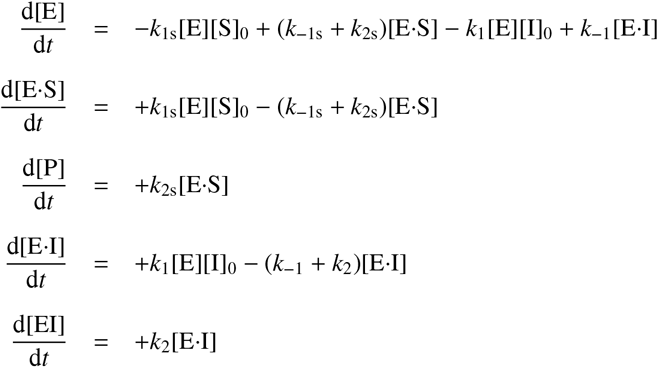

We now invoke the steady-state approximation for the substrate portion of the overall reaction mechanism:

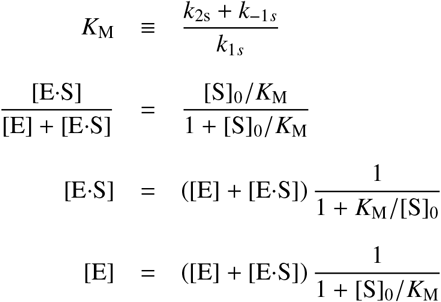

Utilizing the mass balance equation for the enzyme, we obtain the steady state concentrations [E] and [E·S]:

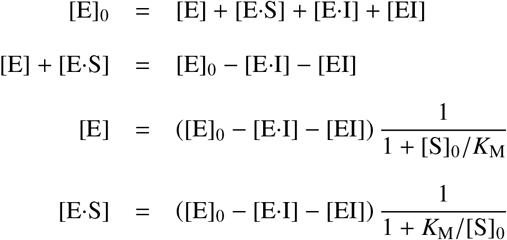

This leads to the reduced system of three simultaneous *linear* differential Eqns (B.8)–(B.10) for three unknowns:

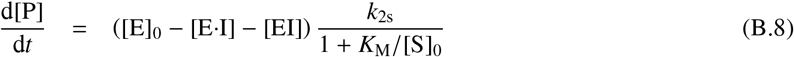

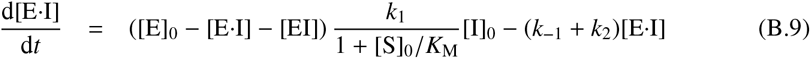

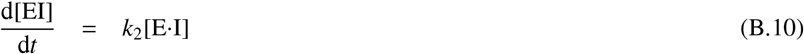

The above system of linear homogeneous ODEs can be solved by the method of Laplace transform. For convenience, we have used the computer-algebra software package Maxima [28]. The integral solution produced by the Maxima Laplace transform algorithm is shown in Eqn (B.11).

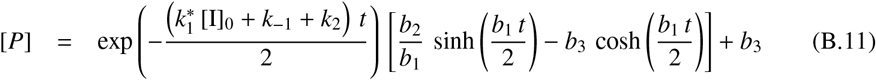

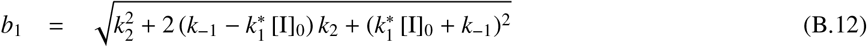

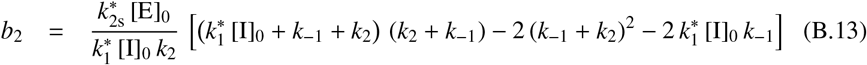

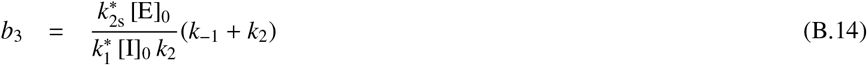

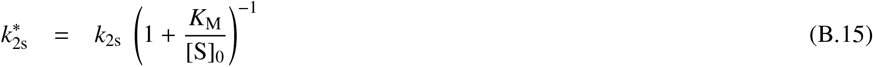

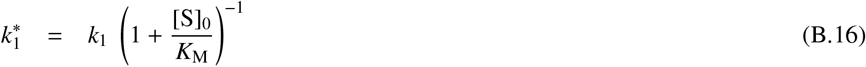

The final algebraic form for the product concentration [P] evolving over time, Eqn (5), was obtained after introducing a number of algebraic simplifications into the integral solution displayed immediately above. In particular, the hyperbolic sine and the hyperbolic cosine functions, which tend to be numerically unstable in actual data fitting, were eliminated by consider their relationship to the relatively numerically stable exponential function:

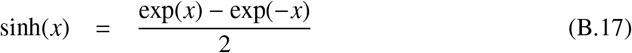

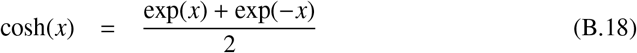

## C. “One-step” kinetics of extremely tight-binding inhibitors

In this Appendix we provide a mathematical proof that a covalent inhibitor characterized by extremely low value of the dissociation constant *k*_−1_ will kinetically follow the one-step inhibition mechanism **C** in *Figure 1*. The proof can be obtained by considering the extreme hypothetical scenario where *k*_−1_ is negligibly small relative to both *k*_1_ × [I]_0_ and *k*_2_. Thus, setting *k*_−1_ = 0 in Eqn (B.9) and integrating the resulting system of linear differential equations Eqns (B.8)–(B.10), using the method of Laplace transform, we obtain for the three state variables a time-dependent solution shown in Eqns (C.1)–(C.3).

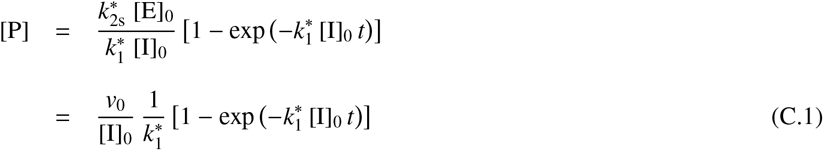

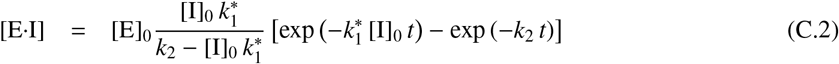

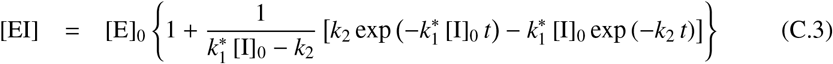

Most importantly, note that the resulting Eqn (C.1) for the product concentration [P] is exactly identical to the one-step kinetic equation Eqn (17). In other words, the assumption that *k*_−1_ = 0, or very nearly so, renders the time-dependence of product concentration entirely insensitive to the value of the inactivation rate constant *k*_2_. In fact, the only rate constant to which [P] is sensitive is the second-order bimolecular association rate constant *k*_1_.

Please note that this conclusion contradicts Cornish-Bowden’s assessment of the steady-state kinetic mechanism **A**. In particular, Cornish-Bowden [11] stated that “if *k*_2_ ≈ *k*_1_ [I]_0_ ≫ *k*_−1_ the two relaxation times [1/*r*_1_ and 1/*r*_2_] defined by [Eqns (9)–(10)] are of similar magnitude and consequently one would observe large deviations from first-order kinetics in any time scale.” This statement is only partially correct, but not in a sense that is relevant to the analysis of continuous assays with real-time monitoring of the product concentration [P]. It is indeed correct to state that if *k*_−1_ happens to negligibly small while *k*2 and *k*_1_ [I]_0_ are of comparable magnitude, “large deviations from first-order kinetics” would be observed *with respect to both enzyme–inhibitor complex concentrations*, E·I and EI, as is shown in in the *double-exponential* equations Eqns (C.2)–(C.3). However, if *k*_−1_ = 0, the *product concentration* [P] changes over time according to the *single-exponential* Eqn (C.1). Thus, with respect to the product concentration changing over time, there are no “deviations” from first-order kinetics and consequently the product concentration change over time exactly as they would do in the case of one-step kinetic mechanism **C**.

See also a detailed analysis of the general kinetic scheme *A* → *B* → *C* presented by Fersht [29, pp.143-144]. This is the simplest case of two consecutive irreversible reactions, governed by rate constants 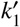 and 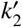 respectively. According to Fersht [29, p. 144, Eqn (4.31)], the concentration of species A, in this case the mole fraction of enzyme that is capable of producing product P in the catalytic cycle, is insensitive to the value of 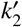 and decays according the *first-order, single-exponential* equation Eqn (C.4).

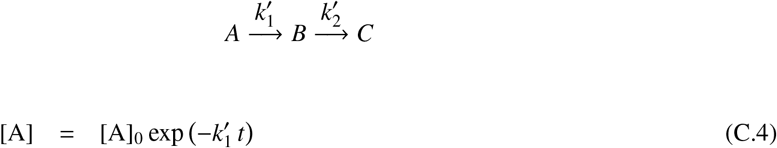

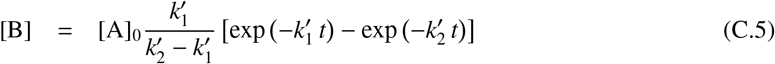

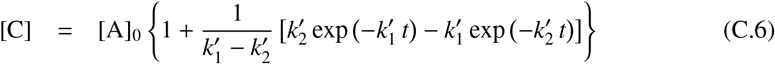

Importantly, because the catalytically competent enzyme fraction decays according to the first-order single-exponential rate law, at *k*_−1_ = 0 the product concentration [P] also follow a (rising) single-exponential defined by Eqn (C.1). Thus, Cornish-Bowden’s prediction [11] that there should be “large deviations from first-order kinetics” is unsupported by detailed theoretical analysis.

## Footnotes

1 RFU stands for relative fluorescence units, but in the more general case it could represent any other appropriate instrument unit such as UV/Vis absorbance units, chromatographic peak areas, radioactive counts, etc.

2 Local optimization means that the optimized parameter is adjustable in the fitting model such that the best-fit value is specific only to a subset of experimental data points, such as in this case each individual progress curve.

