## Supporting Information for "A steady-state algebraic model for the time course of covalent enzyme inhibition"

Petr Kuzmič

*BioKin Ltd., Watertown, Massachusetts, USA*
*http://www.biokin.com*

**2 Microsoft Excel simulation file** **8**

**References** **9**

### 1. DynaFit script file listing

This section lists all DynaFit [1] input script files that were used to generate the present report.

#### *1.1. Structural identifiability analysis*

##### *1.1.1. Simulate ideal data using the algebraic model*

```
[task]
   task = simulate
   data = generic
[parameters]
```

```
   t, Fo, Vo, I
   k1s, km1, k2
[model]
; parameters:
   Vo = 0.0005 ?
   k1s = 0.5 ?       ; k_1^*
   km1 = 0.001 ?     ; k_{-1}
   k2 = 0.01 ?       ; k_2
; model equation:
   gamma = I*k1s + km1 + k2
   alpha = sqrt(gamma*gamma - 4*I*k1s*k2)
   beta = I*k1s*((km1 - k2)/(km1 + k2)) + km1 + k2
   keff = k1s*k2/(km1 + k2)
   a1 = (alpha + beta)/(2*alpha)
   a2 = (alpha - beta)/(2*alpha)
   r1 = (gamma - alpha)/2
   r2 = (gamma + alpha)/2
   F = Fo + Vo/(keff*I)*(1 - a1*exp(-r1*t) - a2*exp(-r2*t))
[data]
   variable t
   mesh from 0 to 3000 step 60
   directory ./TN/2020/02/structural/data/sim-001
   sheet d01-alg.csv
   column  2 | param I = 0.0005| label 0.5 nM
   column  3 | param I = 0.001 | label 1
   column  4 | param I = 0.002 | label 2
   column  5 | param I = 0.004 | label 4
   column  6 | param I = 0.008 | label 8
   column  7 | param I = 0.016 | label 16
[output]
   directory ./TN/2020/02/structural/output/sim-001
[settings]
{Output}
   XAxisLabel = t, sec
   YAxisLabel = F, rfu
[end]
```

#### *1.1.2. Fit simulated data using the corresponding ODE model*

```
[task]
   task = fit
   data = progress
[mechanism]
   E + S <==> E.S    :     k1s    km1s
   E.S ---> E + P    :     k2s
   E + I <==> E.I    :     k1     km1
   E.I ---> EI       :     k2
[constants]
   k1s = 10
   km1s = 9.9
   k2s = 0.1
```

```
   k1 = 1 ?
   km1 = 0.001 ?
   k2 = 0.01 ?
[concentrations]
   E = 0.000001
   S = 1
[responses]
   P = 10000 ?
[data]
   directory ./TN/2020/02/structural/data/sim-001
   sheet d01-alg.csv
   monitor E, E.S, E.I, EI
   column  2 | offset auto ? | conc I = 0.0005| label 0.5 nM
   column  3 | offset auto ? | conc I = 0.001 | label 1
   column  4 | offset auto ? | conc I = 0.002 | label 2
   column  5 | offset auto ? | conc I = 0.004 | label 4
   column  6 | offset auto ? | conc I = 0.008 | label 8
   column  7 | offset auto ? | conc I = 0.016 | label 16
[settings]
{Filter}
   XMin = 1
{Output}
   XAxisLabel = t, sec
   YAxisLabel = F, rfu
[output]
   directory ./TN/2020/02/structural/output/fit-001
[end]
```

#### *1.2. Practical identifiability analysis*

##### *1.2.1. Simulate "noisy" data using the ODE model*

```
[task]
   task = simulate
   data = progress
[mechanism]
   E + S <==> E.S    :     k1s    km1s
   E.S ---> E + P    :     k2s
   E + I <==> E.I    :     k1     km1
   E.I ---> EI       :     k2
[constants]
   k1s = 10
   km1s = 9.9
   k2s = 0.1
   k1 = 0.1 ?
   km1 = 0.001 ?
   k2 = 0.001 ?
[concentrations]
   E = 0.000001
   S = 1
[responses]
   P = 10000
```

```
[data]
   mesh from 0 to 3000 step 60
   error constant 0.5 percent
   directory ./TN/2020/02/practical/data/sim-001
   sheet d01-ode.csv
   monitor E, E.S, E.I, EI
   column  2 | conc I = 0.005| label 5 nM
   column  3 | conc I = 0.01 | label 10
   column  4 | conc I = 0.02 | label 20
   column  5 | conc I = 0.04 | label 40
   column  6 | conc I = 0.08 | label 80
   column  7 | conc I = 0.16 | label 160
[settings]
{Filter}
   XMin = 1
{Output}
   XAxisLabel = t, sec
   YAxisLabel = F, rfu
[output]
   directory ./TN/2020/02/practical/output/sim-001
[end]
```

*1.2.2. Fit simulated data using the corresponding algebraic model*

```
[task]
   task = fit
   data = generic continuous
[parameters]
   t, Fo, Vo, I
   k1s, km1, k2
[model]
; parameters:
   Vo = 0.0005 ?
   k1s = 0.05 ??
   km1 = 0.001 ??
   k2 = 0.001 ??
; model equation:
   gamma = I*k1s + km1 + k2
   alpha = sqrt(gamma*gamma - 4*I*k1s*k2)
   beta = I*k1s*((km1 - k2)/(km1 + k2)) + km1 + k2
   keff = k1s*k2/(km1 + k2)
   a1 = (alpha + beta)/(2*alpha)
   a2 = (alpha - beta)/(2*alpha)
   r1 = (gamma - alpha)/2
   r2 = (gamma + alpha)/2
   F = Fo + Vo/(keff*I)*(1 - a1*exp(-r1*t) - a2*exp(-r2*t))
[data]
   variable t
   directory ./TN/2020/02/practical/data/sim-001
   sheet d01-ode.csv
   column  2 | param Fo = 0 (-1 .. +1) ?, I = 0.005| label 5 nM
   column  3 | param Fo = 0 (-1 .. +1) ?, I = 0.01 | label 10
```

```
   column  4 | param Fo = 0 (-1 .. +1) ?, I = 0.02 | label 20
   column  5 | param Fo = 0 (-1 .. +1) ?, I = 0.04 | label 40
   column  6 | param Fo = 0 (-1 .. +1) ?, I = 0.08 | label 80
   column  7 | param Fo = 0 (-1 .. +1) ?, I = 0.16 | label 160
[output]
   directory ./TN/2020/02/practical/output/fit-001
[settings]
{ConfidenceIntervals}
   SquaresIncreasePercent = 5
{Output}
   XAxisLabel = t, sec
   YAxisLabel = F, rfu
[end]
```

#### *1.3. "One-step" mechanism with high-affinity inhibitor*

##### *1.3.1. Simulate "noisy" data using the algebraic model*

```
[task]
   task = simulate
   data = generic
[parameters]
   t, Fo, Vo, I
   k1s, km1, k2
[model]
; parameters:
   Vo = 0.0005 ?
   k1s = 0.5 ?
   km1 = 0.001 ?
   k2 = 0.01 ?
; model equation:
   gamma = I*k1s + km1 + k2
   alpha = sqrt(gamma*gamma - 4*I*k1s*k2)
   beta = I*k1s*((km1 - k2)/(km1 + k2)) + km1 + k2
   keff = k1s*k2/(km1 + k2)
   a1 = (alpha + beta)/(2*alpha)
   a2 = (alpha - beta)/(2*alpha)
   r1 = (gamma - alpha)/2
   r2 = (gamma + alpha)/2
   F = Fo + Vo/(keff*I)*(1 - a1*exp(-r1*t) - a2*exp(-r2*t))
[data]
   variable t
   mesh from 0 to 3000 step 60
   error constant 0.5 percent
   directory ./TN/2020/02/one-step/data/sim-001
   sheet d01-alg.csv
   column  2 | param I = 0.0005| label 0.5 nM
   column  3 | param I = 0.001 | label 1
   column  4 | param I = 0.002 | label 2
   column  5 | param I = 0.003 | label 3
   column  6 | param I = 0.004 | label 4
   column  7 | param I = 0.005 | label 5
```

```
   column  8 | param I = 0.006 | label 6
   column  9 | param I = 0.008 | label 8
   column 10 | param I = 0.010 | label 10
[output]
   directory ./TN/2020/02/one-step/output/sim-001
[settings]
{Output}
   XAxisLabel = t, sec
   YAxisLabel = F, rfu
[end]
```

*1.3.2. Local fit of simulated progress curves to Kitz–Wilson's equation*

```
[task]
   task = fit
   data = generic
[parameters]
   t, Fo, Vi, kobs
[model]
   F = Fo + (Vi/kobs)*(1 - exp(-kobs*t))
[data]
   variable t
   directory ./TN/2020/02/one-step/data/sim-001
   sheet d01-alg.csv
   column  2 | param Fo = 0 (-1 .. +1) ?, Vi = 0.0005 ?, kobs = 0.001 ? | label 0.5 nM
   column  3 | param Fo = 0 (-1 .. +1) ?, Vi = 0.0005 ?, kobs = 0.001 ? | label 1
   column  4 | param Fo = 0 (-1 .. +1) ?, Vi = 0.0005 ?, kobs = 0.001 ? | label 2
   column  5 | param Fo = 0 (-1 .. +1) ?, Vi = 0.0005 ?, kobs = 0.001 ? | label 3
   column  6 | param Fo = 0 (-1 .. +1) ?, Vi = 0.0005 ?, kobs = 0.001 ? | label 4
   column  7 | param Fo = 0 (-1 .. +1) ?, Vi = 0.0005 ?, kobs = 0.001 ? | label 5
   column  8 | param Fo = 0 (-1 .. +1) ?, Vi = 0.0005 ?, kobs = 0.001 ? | label 6
   column  9 | param Fo = 0 (-1 .. +1) ?, Vi = 0.0005 ?, kobs = 0.001 ? | label 8
   column 10 | param Fo = 0 (-1 .. +1) ?, Vi = 0.0005 ?, kobs = 0.001 ? | label 10
[output]
   directory ./TN/2020/02/one-step/output/fit-001
[settings]
{Output}
   XAxisLabel = t, sec
   YAxisLabel = F, rfu
[end]
```

*1.3.3. Statistical model discrimination analysis of $k_{obs}$ values*

```
;______________________________________________________________________
[task]
   task = fit
   data = generic
   model = two-step ?
[parameters]
   Io, kinact, Ki
[model]
```

```
   kinact = 0.0001 ??
   Ki = 0.01 ??
   kobs = kinact * Io / (Io + Ki)
[data]
   variable Io
   directory ./TN/2020/02/one-step/data/sim-001
   sheet kobs-vs-I.csv
   column 2 error 3
[output]
   directory ./TN/2020/02/one-step/output/fit-002
[settings]
{Output}
   XAxisLabel = [I]_0, {/Symbol m}M
   YAxisLabel = k_{obs}, s^{-1}
   ConfidenceBands = n
   PredictionBands = y
{ConfidenceIntervals}
   LevelPercent = 95
{ModelSelection}
   NestedModels = y
   FCriticalLevelPercent = 95
;______________________________________________________________________
[task]
   task = fit
   data = generic
   model = one-step ?
[parameters]
   Io, keff
[model]
   keff = 0.01 ??
   kobs = keff * Io
[end]
```

### 2. Microsoft Excel simulation file

The Microsoft Excel file BioKin-TN-2020-02-SI2.xls attached to this report contains two sheets named workspace and plots. In order to simulate pseudo-experimental data according to the newly derived two-step steady-state algebraic model, please follow these steps:

1. Open the Excel file BioKin-TN-2020-02-SI2.xls.
2. Switch to the sheet workspace
3. Input numerical values of parameters in the red-bordered rectangle (see *Figure S1*).
4. Set up output points (reaction time in minutes) in column F.
5. Save the modified file.
6. Review the simulated numerical values in columns H–M (see *Figure S2*).
7. Switch to the sheet plots to view the results (see *Figure S3*).

| A | B | C | D | E | F | G |
|---|---|---|---|---|---|---|
| parameters | unit | symbol | value | | time, min | time, s |
| association rate constant for E + S ---> E.S | $\mu M^{-1}\ s^{-1}$ | $k_{1s}$ | 10 | | 0 | 0 |
| dissociation rate constant for E.S ---> E + S | $s^{-1}$ | $k_{-1s}$ | 9.9 | | 2 | 120 |
| "$k_{cat}$" for E.S ---> E + P | $s^{-1}$ | $k_{2s}$ | 0.1 | | 4 | 240 |
| association rate constant for E + I ---> E.I | $\mu M^{-1}\ s^{-1}$ | $k_1$ | 1 | | 6 | 360 |
| dissociation rate constant for E.I ---> E + I | $s^{-1}$ | $k_{-1}$ | 0.001 | | 8 | 480 |
| "$k_{inact}$" for E.I ---> EI | $s^{-1}$ | $k_2$ | 0.01 | | 10 | 600 |
| enzyme concentration | μM | $[E]_0$ | 0.000001 | | 12 | 720 |
| substrate concentration | μM | $[S]_0$ | 1 | | 14 | 840 |
| inhibitor concentration | μM | $[I]_0$ | 0.001 | | 16 | 960 |
| product response | RFU/μM | $r_P$ | 10000 | | 18 | 1080 |
| offset on signal axis | RFU | $F_0$ | 0 | | 20 | 1200 |
| random error | RFU | | 0.01 | | 22 | 1320 |
| | | | | | 24 | 1440 |

**Figure S1:** Microsoft Excel simulation file: Input area.

| F | G | H | I | J | K | L | M |
|---|---|---|---|---|---|---|---|
| | | control progress curve | | | inhibited progress curve | | |
| time, min | time, s | model | error | control | model | error | inhibited |
| 0 | 0 | 0.00E+00 | -1.37E-02 | -1.37E-02 | -1.52E-17 | 1.00E-02 | 1.00E-02 |
| 2 | 120 | 6.00E-02 | 2.95E-03 | 6.30E-02 | 5.83E-02 | 1.13E-02 | 6.95E-02 |
| 4 | 240 | 1.20E-01 | -1.32E-02 | 1.07E-01 | 1.13E-01 | -4.66E-04 | 1.13E-01 |
| 6 | 360 | 1.80E-01 | 9.62E-03 | 1.90E-01 | 1.66E-01 | 1.70E-03 | 1.67E-01 |
| 8 | 480 | 2.40E-01 | -1.62E-03 | 2.38E-01 | 2.15E-01 | -2.07E-03 | 2.13E-01 |
| 10 | 600 | 3.00E-01 | 5.80E-03 | 3.06E-01 | 2.62E-01 | -4.38E-03 | 2.57E-01 |
| 12 | 720 | 3.60E-01 | -1.17E-02 | 3.48E-01 | 3.06E-01 | 2.95E-03 | 3.09E-01 |
| 14 | 840 | 4.20E-01 | -8.72E-03 | 4.11E-01 | 3.48E-01 | 3.12E-03 | 3.51E-01 |
| 16 | 960 | 4.80E-01 | 1.51E-03 | 4.82E-01 | 3.88E-01 | 4.57E-03 | 3.92E-01 |
| 18 | 1080 | 5.40E-01 | 1.66E-02 | 5.57E-01 | 4.25E-01 | 9.39E-03 | 4.35E-01 |
| 20 | 1200 | 6.00E-01 | -1.64E-03 | 5.98E-01 | 4.61E-01 | 2.77E-03 | 4.64E-01 |
| 22 | 1320 | 6.60E-01 | 1.58E-02 | 6.76E-01 | 4.95E-01 | -1.37E-02 | 4.81E-01 |
| 24 | 1440 | 7.20E-01 | 8.45E-03 | 7.28E-01 | 5.27E-01 | 7.09E-03 | 5.34E-01 |

**Figure S2:** Microsoft Excel simulation file: Simulated numerical values.

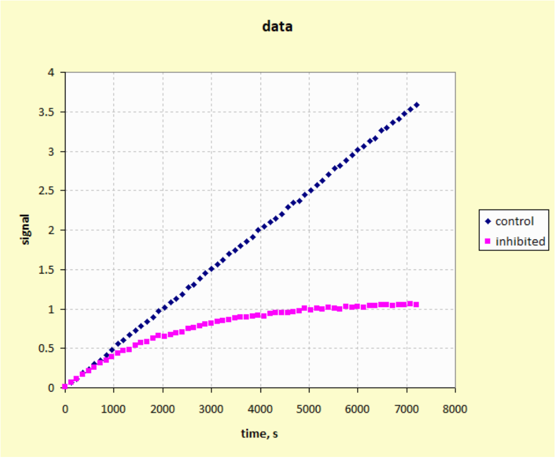

**Figure S3:** Microsoft Excel simulation file: Plot of simulated data.
